# Comparative study of two xanthan gum glycosyltransferases combining AI structure predictions and molecular modeling

**DOI:** 10.64898/2026.03.06.709245

**Authors:** Davide Luciano, Silje Sneve, Gaston Courtade

## Abstract

Xanthan gum is a widely used industrial polysaccharide employed as a thickening and stabilizing agent in food, pharmaceutical, and technological applications. Its biosynthesis involves membrane-associated glycosyltransferases that assemble the repeating unit at the cytoplasmic side of the inner membrane. Among them, GumH and GumI catalyze consecutive reactions using the same donor substrate, guanosine 5’-diphospho-alpha-D-mannose, but with opposite stereoselectivity. Despite their biochemical characterization, structural insights into their catalytic mechanisms and membrane interactions remain limited, hindering a detailed understanding of their function and future engineering efforts. In this work, we combined artificial intelligence-based structure prediction with atomistic molecular dynamics simulations to investigate the structural organization and substrate-binding modes of GumH (family GT4) and GumI (family GT94). The predicted apo structures exhibit a conserved GT-B fold but differ in interdomain flexibility and membrane-anchoring strategies. GumH displays a more structured interdomain linker and a defined clamp-like region in the acceptor-binding domain, consistent with stable membrane interaction, whereas GumI shows a more flexible linker and an open groove architecture. Modeling of the donor-bound complexes reveals distinct substrate-binding modes. In GumH, it adopts a geometry consistent with its retaining stereochemical outcome, positioning the sugar close to the conserved catalytic residue. In contrast, GumI exhibits a different donor orientation, lacking a clearly positioned catalytic base near the reactive center, suggesting a substrate-assisted catalytic mechanism. Although the predicted ternary complexes show limited stability in our simulations, they provide chemically reasonable conformations and offer structural insights into substrate recognition, membrane association, and stereochemical control in these two glycosyltransferase families.

**Significance statement:** Xanthan gum is an industrially important polysaccharide widely used in food and other technological products. Although several enzymes in its biosynthetic pathway have been studied, structural information remains limited. Using AI-based structure predictions and molecular simulations, we revealed how these enzymes sit in the membrane and bind sugar substrates. These structural insights clarify xanthan biosynthesis and could help improve or engineer its production.

## 1 Introduction

Exopolysaccharides (EPS) are biologically abundant molecules in the extracellular matrix of many microorganisms. EPS are composed of long chains of monosaccharides linked by glycosidic bonds and have been widely studied for both their industrial applications and their role in pathogenic activity [1]. Considerable research has focused on elucidating the biochemical pathways involved in their biosynthesis [2, 3].

EPS are typically synthesized through various strategies, which in some cases involve glycosyltransferases (GTs), enzymes that catalyze glycosidic bond formation and assemble the repeating units of the polymer [4]. These enzymes generally bind two substrates: (i) an acceptor substrate, which in the case of EPS is the growing repeating unit, and (ii) a donor substrate, which is most often a nucleotide sugar. Within EPS biosynthetic pathways, these enzymes are usually found in close association with the membrane. Despite active research in this field, the molecular mechanisms that govern GT activity and selectivity remain largely unknown.

Among the many EPS, xanthan gum is among the most widely used in industry [5]. Developing pipelines to efficiently modify this polysaccharide is an important step toward the production of novel biomaterials. The repeating unit of xanthan gum consists of a cellulosic backbone composed of two *β*-glucose units connected by a *β*-1,4 glycosidic bond, and a trisaccharide side chain composed of *α*-mannose, *β*-glucuronate, and *β*-mannose. This side chain can be further modified by non-carbohydrate substitutions, most notably pyruvate and acetyl groups on the first and last mannose residues [6]. The sequence of monosaccharides in xanthan gum is highly conserved, indicating the high substrate specificity of the enzymes involved in its biosynthesis. However, the biochemical mechanisms underlying this selectivity by the GTs responsible for xanthan biosynthesis remain unclear.

The biosynthetic pathway of xanthan gum can be conceptually divided into two stages [7]: (i) assembly of the lipid-linked repeating unit by a series of cytosolic or membrane-associated GTs, and (ii) polymerization, export, and release of the mature polysaccharide into the extracellular matrix. Although many components of this pathway remain poorly characterized, GTs are particularly attractive targets for investigation because modifications in their specificity directly translate into changes in the chemical composition of the final biomaterial.

Within the side-chain biosynthetic module (Fig. 1), three GTs play a central role. GumH [8] catalyzes the addition of an *α*-mannose residue to the growing repeating unit, generating the acceptor substrate for GumK[9], which subsequently transfers a *β*-glucuronate moiety. Finally, GumI[10] completes the side chain by adding a terminal *β*-mannose residue. Notably, GumH and GumI both utilize *α*-mannose-GDP as donor substrate, yet catalyze reactions with opposite stereochemical outcomes: GumH is a retaining GT, whereas GumI is an inverting GT. In addition to this difference, these enzymes belong to distinct CAZy [11] families—GT4 for GumH and GT94 for GumI—reflecting differences in their evolutionary origins and overall folds.

**Figure 1.**
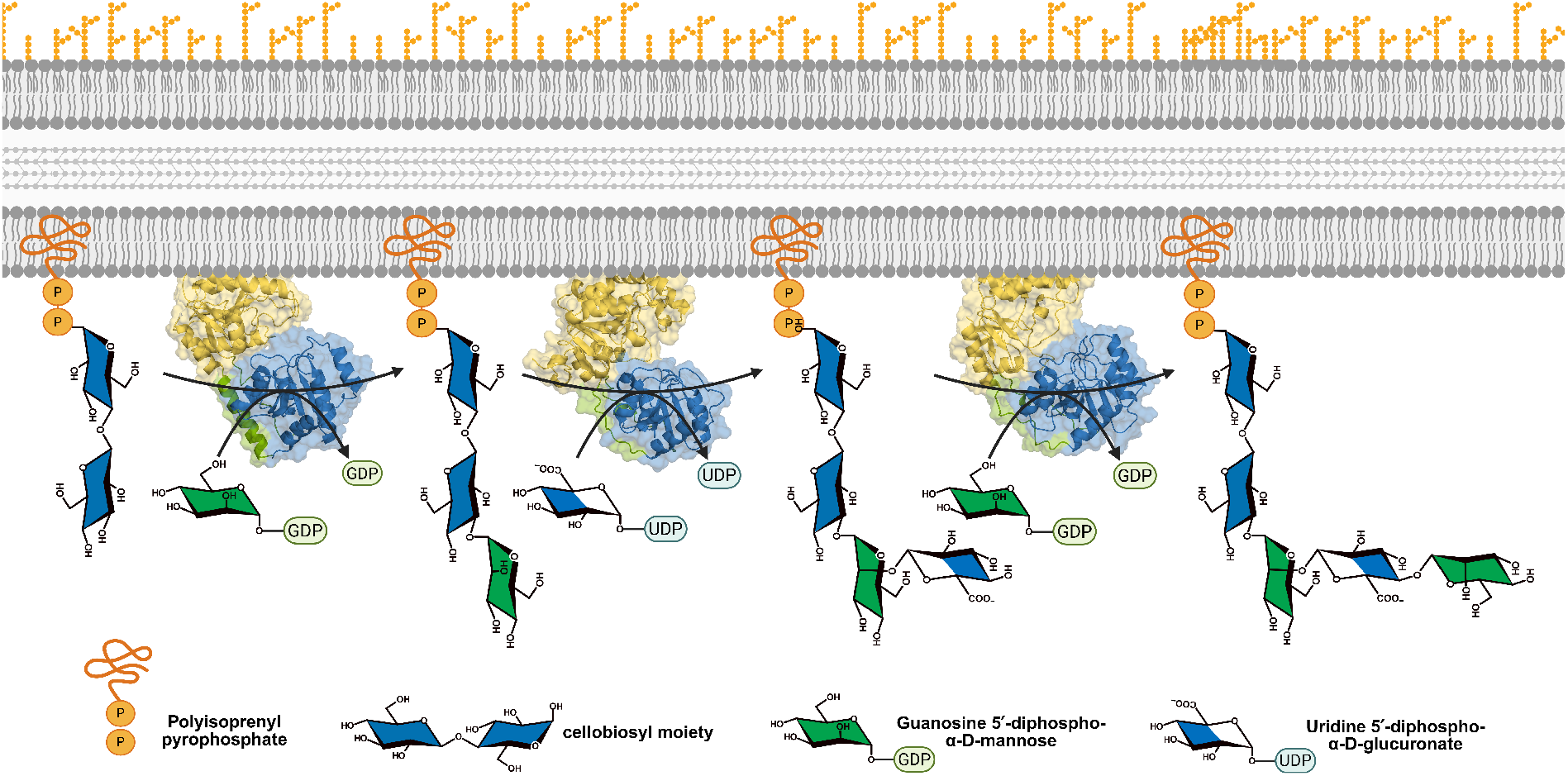
Biosynthetic pathway of xanthan gum side chains and the three glycosyltransferases involved in the process. The side chain of xanthan gum is assembled stepwise by three enzymes. First, GumH (CAZy GT4) transfers an *α*-mannose from GDP-*α*-mannose to the cellobiosyl moiety biosynthesized in the upstream steps. Then, GumK (CAZy GT70) adds a *β*-glucuronate from UDP-*β*-glucuronate to the product formed by GumH. Finally, GumI (CAZy GT94) completes the side chain by transferring a *β*-mannose from GDP-*α*-mannose to the product of GumK. The final product corresponds to the complete repeating unit, which will be polymerized in the downstream steps of the pathway. The assembly happens at the membrane, where the growing structure is anchored through the lipid carrier undecaprenyl pyrophosphate.

The enzymatic activities of GumH and GumI have been biochemically characterized [8, 10, 12], but their experimentally-determined high-resolution structures remain unknown. This lack of structural insight represents a major limitation for understanding their catalytic mechanisms and hampers rational efforts to engineer xanthan gum biosynthesis. In contrast, GumK has been studied more extensively [9, 13], and we recently reported a computational characterization of this enzyme, supported by the available crystal structures of GumK in its *apo* form and in complex with UDP [14].

Recent advances in artificial-intelligence–based structure prediction [15], most notably AlphaFold [16] and Boltz [17], have transformed structural biology by enabling accurate modeling of protein folds and, in some cases, protein–ligand complexes [18]. However, these methods have significant limitations when applied to carbohydrate substrates, particularly in reliably distinguishing between different chiral configurations and glycosidic linkages [19].

In this work, we combine AI-based structure prediction with atomistic molecular dynamics simulations to investigate GumH and GumI in complex with their native substrates, providing structural insights into substrate recognition and catalytic organization within two distinct CAZy families.

Our results offer a framework for interpreting existing biochemical data and for future mutagenesis and enzyme engineering, and help identify generalizable structure–function principles across GT families.

## 2 Results

### 2.1 The apo forms of GumH and GumI show distinct domains, dynamics, and membrane anchoring strategies

GumH (UniProt: Q56774) and GumI (UniProt: Q56775) are GT-B enzymes belonging to CAZy families GT4 and GT94 [11]. GT-B proteins adopt two Rossmann-like domains, N- and C-terminal, connected by a flexible linker [20], enabling inter-domain twisting and bending motions often required for catalysis. As in other GT-B enzymes, the domains bind distinct substrates: the N-domain binds the membrane-anchored oligosaccharide acceptor, whereas the C-domain binds the donor *α*-GDP-mannose [10, 12].

Both proteins interact with membranes, but in different ways. GumI is proposed to be a monotopic membrane protein [10], while GumH is described as cytoplasmic [4], suggesting membrane interaction upon acceptor binding. Without structural data, the molecular basis and functional consequences of membrane association remain unclear.

To investigate the apo forms, we considered two conformations: model0, the highest-confidence Boltz prediction without substrates, and conf0, a more closed inter-domain state derived from the predicted ternary complex. conf0 serves as a reference for inter-domain dynamics.

Boltz, like AlphaFold3, combines template information and MSA within a generative framework. The best GumH model (model0) is shown in Figure 2A, colored by pLDDT. Scores for all ten models are reported in Figure S1A. Across the ensemble, the average pLDDT exceeds 0.9, indicating high local confidence.

**Figure 2.**
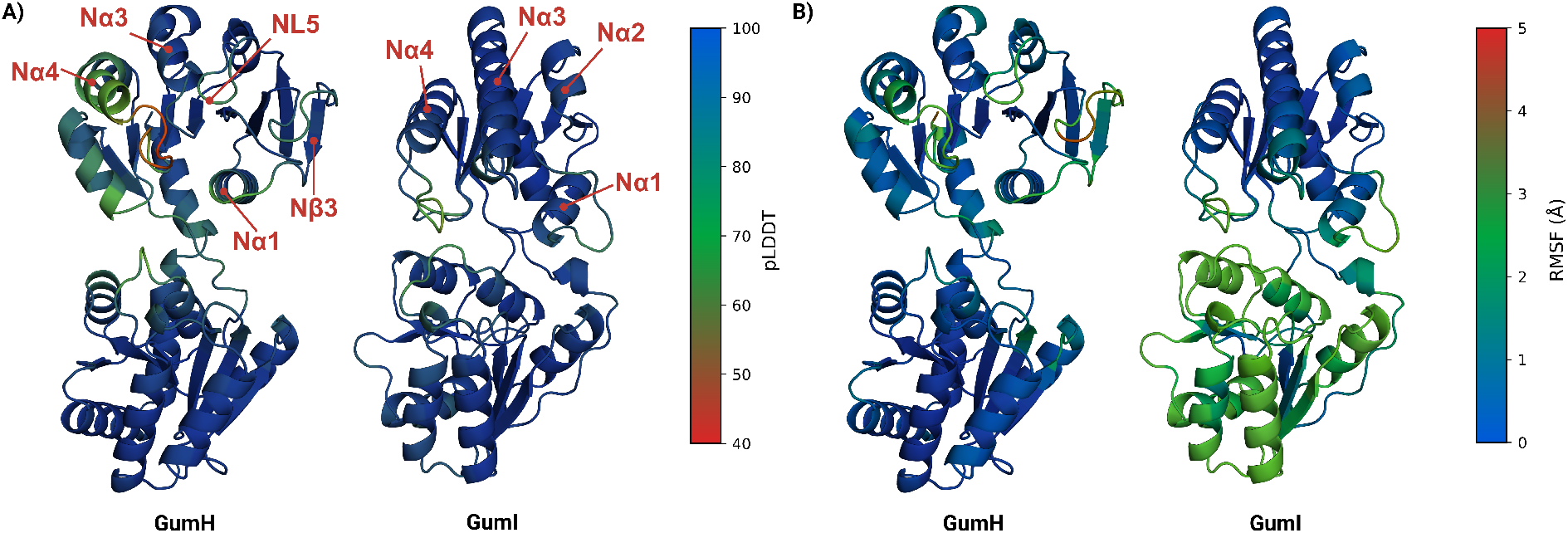
A) Boltz predictions of the apo form for GumH and GumI. The structures are colored based on the pLDDT. The key secondary structures discussed in the main text are also labeled. B) The predicted conformations are colored based on the root mean squared fluctuation of the alpha carbons for GumH and GumI. The RMSF values are averages across simulations of the apo protein anchored to the membrane, the predicted apo form, and the three closed-form replicas.

In the acceptor-binding domain, a loop connected to helix *Nα*4 shows low pLDDT values (Fig. 2A). Given the high sequence coverage in this region (Fig. S2A), this likely reflects intrinsic flexibility. Helix N*α*4 displays pLDDT >70 except at its junction with the loop, indicating reliable backbone but uncertain side chains [16, 21]. Despite high local confidence, PAE and pTM scores suggest flexibility in the loop and inter-domain arrangement. Inter-domain distances across the ten models confirm sampling of multiple conformations with varying domain bending.

Compared to conf0, all predicted folds are more open, with model0 being the most open (Fig. S3A). The structure includes a hydrophobic N*α*4 helix (Fig. 2A) containing serine, threonine, and positively charged residues (Fig. S4), typical of membrane-associated helices [22]. However, residues commonly involved in monotopic anchoring, such as tryptophan, are absent.

The Boltz prediction of GumI also displays a canonical GT-B architecture (Fig. 2A). Average scores across the ten models indicate stable intradomain folds, with some interdomain flexibility suggested by PAE (Fig. S5A). Inter-domain distances remain larger than in conf0 (Fig. S3B). Residue-wise pLDDT values (Figure 2A) indicate higher overall confidence than in GumH. Low-confidence regions do not correspond to defined secondary structures and, given the high sequence coverage (Fig. S2B), likely reflect intrinsic flexibility.

GumI has been proposed as a monotopic membrane protein [10]. Consistently, the predicted structure shows an extended hydrophobic surface patch involving helices N*α*2, N*α*3, and N*α*4 (Fig. 2A). Helix N*α*4 contains two exposed tryptophans, Trp100 and Trp108, which contribute to this patch and are arranged similarly to those reported for GumK [14].

To probe the dynamics of the predicted folds of GumH and GumI, we performed an anisotropic elastic network model (ANM) analysis [23] using ProDy [24]. In ANM, backbone fluctuations are described within a harmonic approximation by modeling the protein as an elastic network of C_*α*_ atoms connected within a distance cutoff. The collective motions of this network are represented by vectors, known as normal modes, which define the direction and amplitude of concerted displacements. These modes are ranked by frequency, and the lowest-frequency ones correspond to large-scale conformational changes that are often functionally relevant. The analysis was carried out on model0 from the Boltz predictions and on the closed reference conformation conf0 derived from the predicted ternary complexes.

To approximate native conditions, we included an elastic membrane model [25]. Protein orientation was predicted with OPM [26] assuming a Gram-negative inner membrane. The resulting orientations (Fig. S6A,B) agree with the proposed membrane-interacting regions.

For GumH, model0 shows dominant side-bending motions (Fig. S7A). In conf0, the first mode is twisting, followed by side and frontal bending (Fig. 3A).

**Figure 3.**
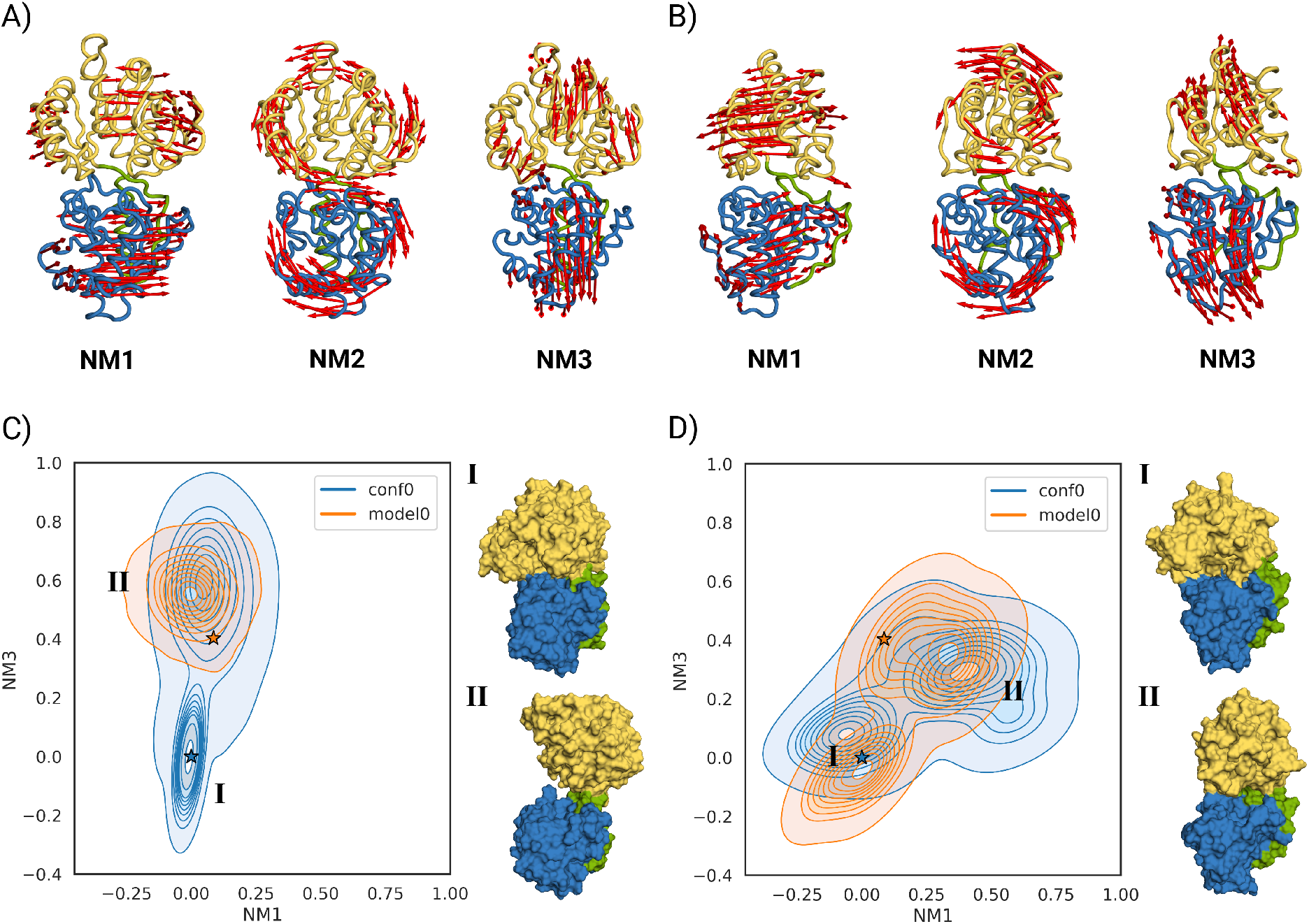
A) First three normal modes computed starting from the closed conformation of GumH. The first normal mode (NM1) involves twisting of the two domains, and the second (NM2) involves side bending that causes the two domains to rotate in opposite directions. The third (NM3) is the frontal bending, which opens the two domains. B) First three normal modes computed starting from the closed conformation of GumI. The first normal mode (NM1) involves twisting of the two domains, and the second (NM2) involves side bending that causes the two domains to rotate in opposite directions. The third (NM3) is the frontal bending, which opens the two domains. C) Ensemble of conformations sampled during the unbiased MD simulations of the predicted apo form of GumH model0 (orange) and the predicted closed conformation conf0 (blue). Two populations are clearly visible: a closed conformation (state I) and an open conformation (state II). The stars indicate the starting conformations. On the side, there are two structures that represent the respective states. D) Ensemble of conformations sampled during the unbiased MD simulations of the predicted apo form of GumI model0 (orange) and the predicted closed conformation conf0 (blue). Two populations are clearly visible: a closed conformation (state I) and a twisted semi-open conformation (state II). The stars indicate the starting conformations. On the side, two structures represent the respective states.

For GumI, model0 exhibits twisting and bending as the first two modes, followed by side bending (Fig. S7B). In conf0, the second and third modes are inverted (Fig. 3B). These differences reflect distinct elastic network topologies (Fig. S7C,D) due to domain orientation. Since conf0 shows more inter-domain contacts, its first two normal modes were used as references.

We used ClustENMD [27] to apply a deforming force to the proteins. We deformed conf0 along the first three modes. Thirty generations were generated with a target RMSD of 1 Å, and the process was repeated three times. Generated conformations were projected onto the first and third modes (Fig. 4A,C).

**Figure 4.**
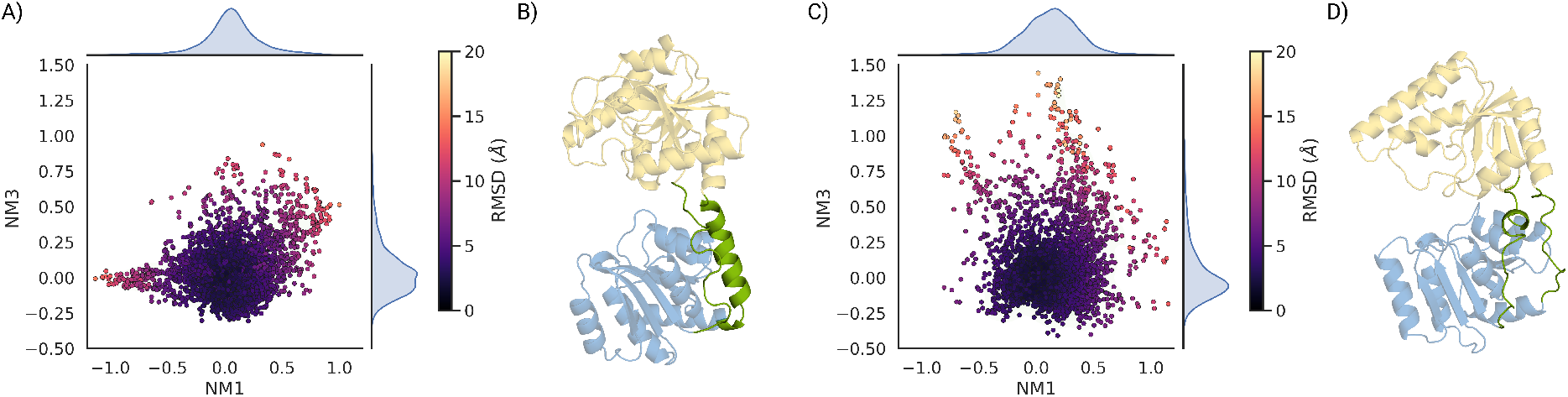
A) ClustENMD ensemble of GumH conformations starting from the closed form and projected on the first (NM1) and third (NM3) normal modes. The points are colored by the RMSD of each conformation relative to the starting conformation, which is the origin of the plot. B) linker region of GumH between the acceptor-binding domain and the donor-binding domain, which shows an extra halpa helix that makes the opening of the protein more difficult. C) ClustENMD ensemble of GumI conformations starting from the closed form and projected on the first (NM1) and third (NM3) normal modes. The points are colored by the RMSD of each conformation relative to the starting conformation, which is the origin of the plot. D) Linker region of GumI between the acceptor-binding domain and the donor-binding domain, which shows a less structured region, making the protein more prone to deformation and overtwisting.

GumH conformations remained close to conf0, with limited RMSD deviation. In contrast, GumI sampled more open states. The largest RMSD corresponds to an overtwisted conformation in which the acceptor-binding domain rotates 180° relative to the starting position (Fig. S8).

To investigate membrane-anchored behavior, model0 and conf0 from GumH and GumI were inserted into a lipid bilayer built with CHARMM-GUI to resemble the Xanthomonas inner membrane [28]. OPM orientations (Fig. S6A,B) were used. All simulations ran for 1 µs.

For GumH, the apo model0 remains near its starting conformation (Fig. 3C). conf0 opens and converges toward the same ensemble, confirming that the predicted apo state corresponds to an open conformation. Opening mainly involves frontal bending rather than twisting.

RMSF analysis (Fig. 2B) identifies two flexible regions in the acceptor-binding domain: residues 118 to 126 and 44 to 51. The first corresponds to the lowest-confidence region. The second displays the highest flexibility and partial unfolding, propagating to N*β*3 strand detachment.

The simulations show that the region between helix N*α*4 and loop NL5 (Fig. 2A) behaves as a clamp. Its state is described by the C*α* distance between Tyr74 and Lys131 (*d*_clamp_; Fig. 5). model0 samples fully open (state IV) and semi-open (state III) states, depending on phospholipid type. The semi-open state interacts with phosphatidylglycerol, whereas the fully open state is more frequent with phosphatidylethanolamine. conf0 samples the semi-open state (III) and a fully closed state (I), which does not interact with phospholipids.

**Figure 5.**
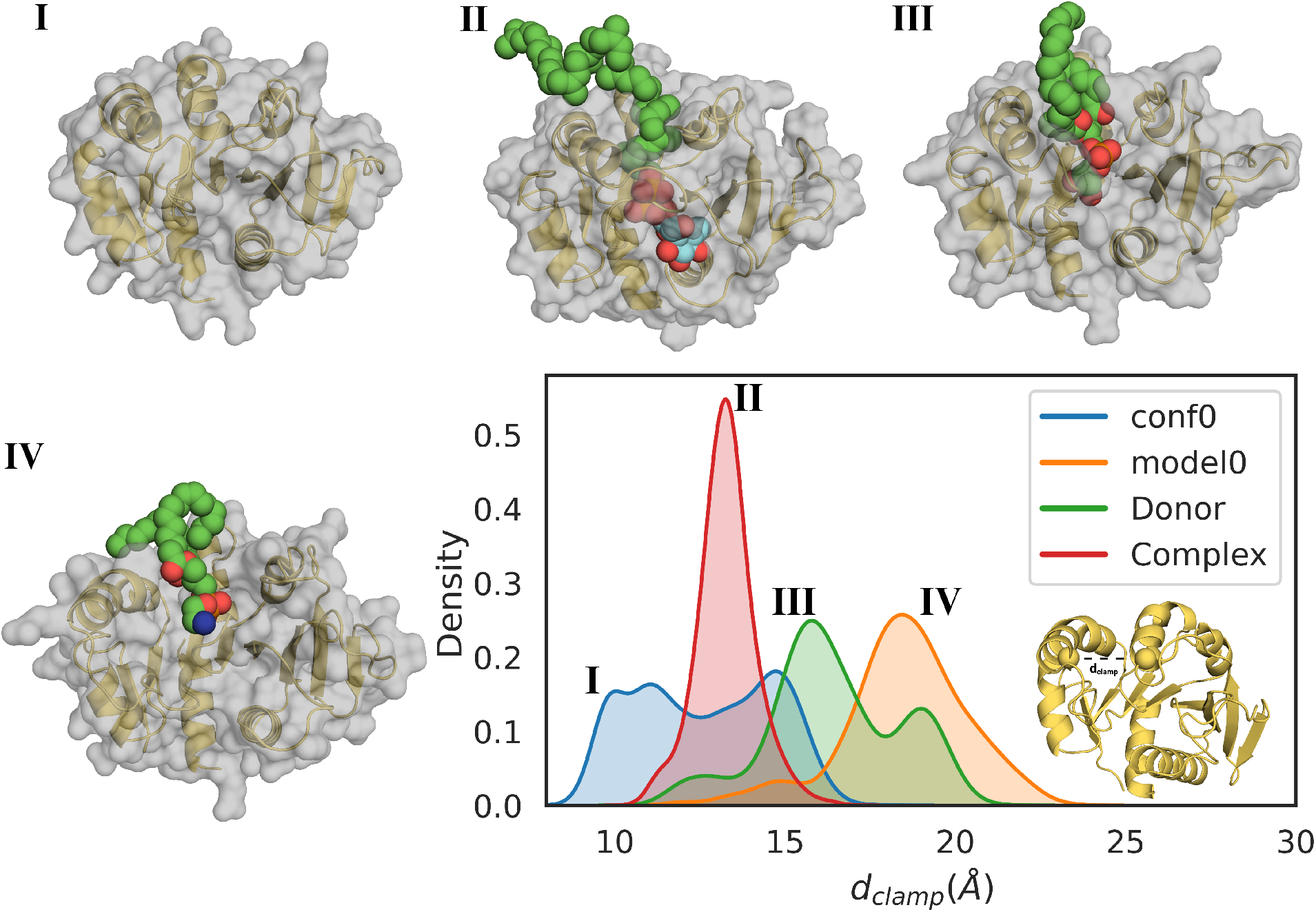
Structures of ternary complexes of GumH and distance distribution of the clamp region, *d*_*clamp*_. This distance is used to describe the different states of the clamp motion in the acceptor-binding domain of GumH. The different colors represent different unbiased molecular dynamics simulations, starting from four different conformations: apo protein in the closed conformation, conf0 (blue), predicted apo protein in the open conformation, model0 (orange), GumH in complex with the donor substrate (green), and GumH in complex with both the donor and acceptor substrates (red). The structures numbered I-IV represent different states identified in the simulations: a fully closed conformation represented (state I), a conformation in which the calmp binds the lipidic tail of the acceptor substrate (sate II), a semi-open conformation (state III) usually observed in complex with phosphatidylglycerol, and a fully open conformation (state IV), which was observed intercting with positively charged phospholipids.

Overall, helices N*α*2–N*α*4 (Fig. 2A) form a dynamic clamp capable of adopting multiple conformations and interacting with distinct phospholipids.

For GumI, MD simulations starting from model0 and conf0 were projected onto the first and third modes (Fig. 3D). model0 samples both open and closed states; in one replica, it adopts a more closed conformation. conf0 progressively opens and converges toward the model0 ensemble. However, unbiased MD does not sample fully open states as observed with ClustENMD.

RMSF analysis (Fig. 2B) shows that GumI maintains a stable fold without unfolding. The donor-binding domain is more flexible than the acceptor-binding domain, and low pLDDT regions (Fig. 2A) correspond to flexible segments.

During simulation, phospholipid extraction events occur between helices N*α*2 and N*α*3 (Fig. 2A). Tyr71, located in a flexible loop, facilitates these interactions. This region likely represents the binding site of the lipidic acceptor carrier and contains positively charged residues that can interact with its pyrophosphate group. Unlike GumH, GumI lacks a defined clamp motif and instead presents a rigid groove.

In summary, Boltz predicts relatively open apo conformations for both proteins. Although their folds are similar, GumH and GumI exhibit distinct membrane-anchored dynamics. Both interact stably with the membrane and bind phospholipids in the acceptor-binding region. In GumI, this region is less structured, leading to faster phospholipid exchange than in GumH, whose fold shows a more organized interaction surface.

### 2.2 GumH and GumI bind the donor substrate differently

To investigate the molecular basis of the different stereoselectivities of GumH and GumI, we predicted donor–substrate complexes using Boltz and analyzed their structural features and dynamics.

Boltz supports ligand input via SMILES or CCD codes; the latter is recommended to preserve crystallographic geometry [17]. GDP-Man (CCD code GDD), well characterized in the PDB, was therefore used.

As in the apo predictions, ten models were generated for each donor-bound complex. The folding is consistent with the apo-predictions, and GDP-Man binds the C-terminal domain.

For GumH, the predicted complexes show stable confidence scores across models, with high reliability for both the fold and the protein–ligand complex (Fig. S1B). The only weaker metric is the ipDE score, indicating uncertainty in ligand positioning.

Compared to the apo state, donor-bound models show more consistent inter-domain distances, closer to conf0, indicating a more closed arrangement (Fig. S3C). Residue-wise pLDDT values (Fig. S9A) increase overall relative to the apo form, although the loop near helix N*α*4 (Fig. 2A) remains low-confidence, suggesting persistent flexibility.

The first predicted conformation was selected for further analysis (Fig. S9A). In this model, GDP-Man interacts with Tyr181 and Glu287 via ribose hydroxyl groups, while the purine ring contacts the backbone of Pro257, with Ile14 stacking against it. The diphosphate forms polar interactions with Arg198 and Lys203 and is oriented approximately perpendicular to the purine plane, further stabilized by the N*α*1 helix backbone (Fig. 2A). The mannose moiety bridges both domains, interacting with Asn174 and His118 in the acceptor domain and residues Glu279–Ile283 in the donor domain. This segment includes the conserved E*X*_7_E motif of GT4 enzymes [29]; Glu279 is positioned near the anomeric carbon, whereas Glu287 primarily contacts the ribose.

The GumH donor-bound complex was inserted into a Xanthomonas-like bilayer using OPM orientation and CHARMM-GUI, following the apo protocol. Three 500 ns replicas were performed.

The sampled conformations were projected onto the first (twisting) and third (bending) normal modes (Fig. 6A). Compared to the apo form, the protein remains more compact and shows enhanced twisting. Despite this closed conformation, native contacts between substrate and protein decrease to 40–50%, mainly due to reorientation of the mannose moiety, while the guanosine remains stably bound (Fig. S8A). As in the apo simulations, the region near N*β*3 remains unstable, suggesting local inaccuracies in the predicted fold.

**Figure 6.**
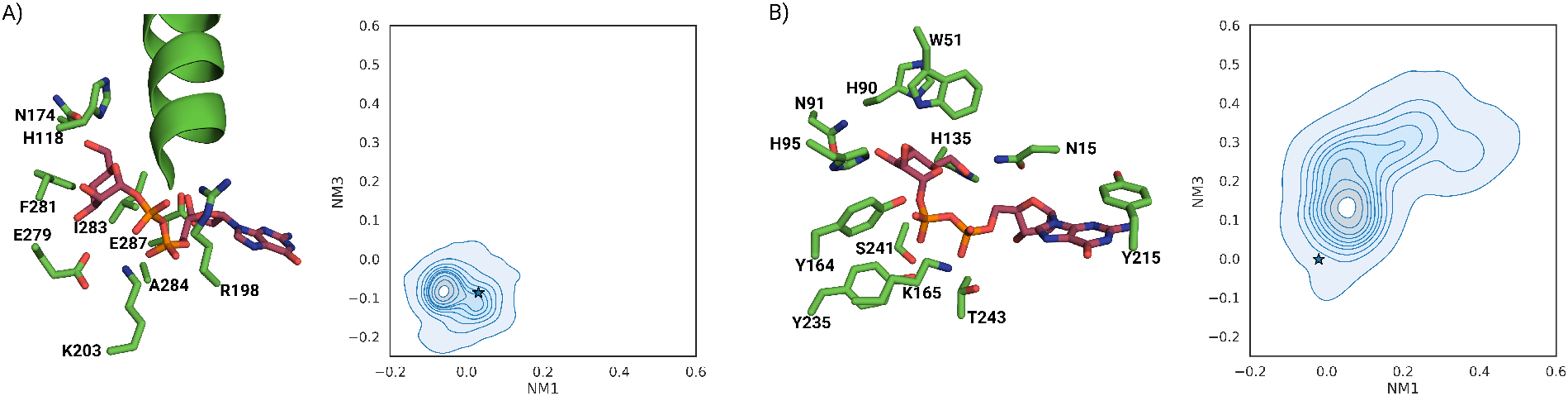
A) On the left, the equilibrated structure of the model0 fold predicted for the GumH–donor complex. On the right, the ensemble of protein conformations sampled during the three unbiased molecular simulations is projected onto the first normal mode (domain twisting) and the third normal mode (domain bending).The stars indicate the starting conformations. B) On the left, the equilibrated model0 conformation predicted for the GumI–donor complex. On the right, the ensemble of conformations sampled during the three unbiased molecular simulations is projected onto the first normal mode (domain twisting) and the third normal mode (domain bending). The stars indicate the starting conformations.

To obtain a more robust statistical characterization of the clamping region in the acceptor-binding domain, we analyzed the conformational states sampled during the simulations. All replicas populate the same clamp conformations (states III and IV) observed in the apo simulations (green distributions in Fig. 5), further supporting the functional role of this region in acceptor binding.

For GumI, donor-bound predictions follow a similar trend. The global fold is predicted with high confidence, while ipDE indicates interface uncertainty. Inter-domain distances are more consistent than in the apo form but remain shifted toward more open conformations relative to conf0 (Fig. S3D).

Residue-wise pLDDT analysis (Fig. S9B) indicates greater confidence than in the apo form. In the predicted binding mode, the guanosine moiety stacks with Tyr215 and forms hydrogen bonds via the ribose C3 hydroxyl with Tyr138 and the backbone of Tyr215 and Val216. The diphosphate interacts with Thr243, Lys165, Tyr235, Tyr164, and Ser241 and adopts a coplanar orientation relative to the guanosine ring, distinct from that in GumH. The mannose moiety interacts extensively with the acceptor domain, forming polar contacts with Trp51, Asn91, His90, His135, and Asn15. A *π*–*π* stacking interaction between His95 and Tyr164 stabilizes the closed inter-domain state. The equilibrated first model structure is shown in Fig. 6B.

As for GumH, three 500 ns unbiased MD simulations were performed with membrane anchoring via OPM and CHARMM-GUI, totaling 1.5 µs. In two replicas, the protein opens along a pathway similar to the apo form, involving both bending and twisting (Fig. 6B). The mannose moiety is unstable and flips during simulation, reducing native contacts to 20%, whereas the guanosine remains stably bound (Fig. S10B).

As observed in the apo state, the groove between helices N*α*2 and N*α*3 (Fig. 2A) exchanges phospholipids, reinforcing its proposed role in acceptor binding.

Overall, Boltz predicts a more closed inter-domain arrangement for both enzymes in the donor-bound state, with distinct binding modes for GumH and GumI. Although fold confidence increases relative to the apo form, the donor-bound complexes do not remain fully stable during solution-phase simulations.

### 2.3 Ternary complexes reveal a dynamic clamp that stabilizes the acceptor substrate

Having characterized donor-membrane interactions, we built full-complex models of GumH and GumI using Boltz. We predicted ternary complexes containing both substrates and product-bound states to obtain suitable starting structures for modeling reactive configurations.

Although both enzymes bind GDP-Man, they act on distinct but structurally related acceptors that are sequential intermediates of the same pathway. GumH transfers mannose to the C3 position of glucose-*β*-1,4-glucose-*α*-1-UndPP via an inverting mechanism. GumI acts on glucuronate-*β*-1,2-mannose-*α*-1,3-glucose-*β*-1,4-glucose-*α*-1-UndPP and transfers mannose to C4 of the glucuronate unit, also via inversion (Fig. 1).

For GumH, both ternary complexes show low confidence at the substrate interface across the ten predicted models (Fig. S1C,D). Residue-wise pLDDT analysis (Fig. S9C) indicates that the acceptor and product are the least reliable regions, whereas the protein fold is predicted with higher confidence. pLDDT values are particularly low for the lipidic tail, while the saccharidic portion is better resolved. In model 0 of both complexes, the acceptor ring is distorted; in the substrate-bound state, the wrong hydroxyl group (C2 instead of C3) is positioned near the donor anomeric carbon.

The lipidic tail binds in the same region identified in apo and donor-bound simulations, supporting the clamp hypothesis in the acceptor-binding domain. The donor position resembles that observed in the donor-only complex. In the product complex, the transferred mannose occupies a position similar to that of the donor mannose, suggesting positional bias in the prediction.

Because the substrate-bound model is improperly docked and model0 of the product complex shows ring distortion, we selected model 2 from the product predictions for refinement. All ten product-bound models display comparable confidence (Fig. S1D), and model 2 lacks major deformations.

After restoring substrate connectivity from model 2 and optimizing geometry with YASARA, the complex was inserted into a Xanthomonas-like membrane using the same protocol described above. Three independent 500 ns simulations were performed.

The equilibrated complex is shown in Fig. 7. The acceptor is oriented such that the C3 and C4 hydroxyl groups lie at 3.3 Å and 3.7 Å from the donor anomeric carbon, respectively. The C3 hydroxyl is closer to the diphosphate moiety, with an angle of 72.8° between the donor O1–C1 vector and the acceptor O3 atom, compared to 106.2° for O4. This geometry favors interaction between the *α*-phosphate and O3, positioning O3 on the same side of the phosphate group, consistent with a retaining mechanism. The orientation is stabilized by interactions with Glu18, Arg8, and His123 in the N-domain, and Arg198 and Tyr231 in the donor-binding domain.

**Figure 7.**
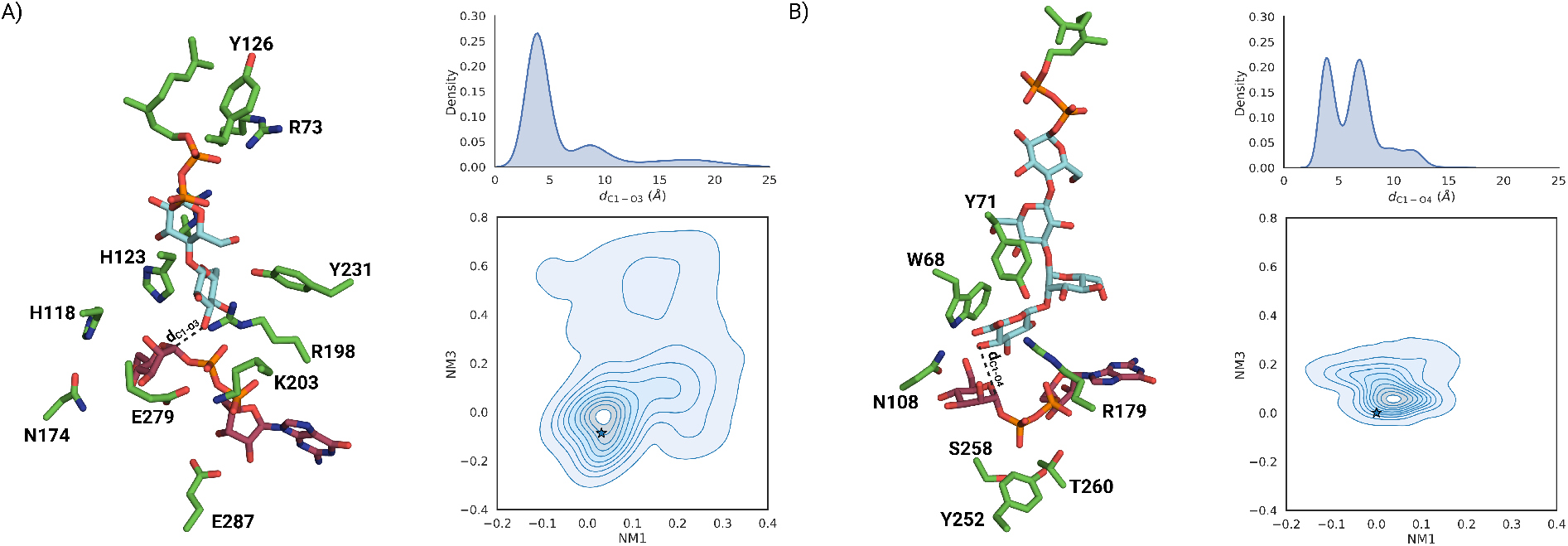
A) On the left, the equilibrated full-complex conformation of GumH predicted using Boltz is shown. The cyan rings represent the acceptor substrate, and the purple ligand represents the donor substrate (GDP-mannose). On the top right, the distribution of the reactive distance between the anomeric carbon of the donor substrate and the O3 of the acceptor substrate (*d*_*C*1−*O*3_) sampled during the simulation is reported. In the bottom-right panel, the protein conformations sampled during the three full-complex simulations are projected onto the space defined by the first and third normal modes. The stars indicate the starting conformations. B) On the left, the equilibrated full-complex conformation of GumI predicted using Boltz is shown. The cyan rings represent the acceptor substrate, and the purple ligand represents the donor substrate (GDP-mannose). On the top right, the distribution of the reactive distance, defined as the distance between the O4 of the acceptor substrate and the anomeric carbon of the donor substrate, is shown. In the bottom-right panel, the protein conformations sampled during the simulations are projected onto the space defined by the first and third normal modes. The stars indicate the starting conformations.

The complex remains stable for over 100 ns, as shown by the C1–O3 reactive distance (Fig. S10C), whose main population is compatible with a reactive state (Fig. 7A). In two of three replicas, the protein stays closed, while twisting increases, as shown by projection onto the first and third normal modes (Fig. 7A).

Monitoring the clamping distance (red distribution in Fig. 5) reveals a new intermediate state (state II) between the closed (I) and semi-open (III) conformations. This behavior likely reflects the presence of a single lipidic tail, which induces a more open clamp than phospholipid double chains. The tail remains stabilized by Tyr126 and Arg73 (Fig. 7A), further supporting the clamp’s role in stabilizing the reactive configuration.

For GumI, ternary complex predictions show a similar trend: the protein fold is consistent with previous models, whereas ligand placement is less confident (Fig. S5C,D), as supported by residue-wise pLDDT analysis (Fig. S6). The absence of a lipidic carrier increases overall confidence compared to GumH.

The first model of the substrate-bound prediction shows deformation of the glucuronate unit and incorrect C2 stereochemistry in the donor. We therefore selected the first prediction from the product-bound complex, which lacks critical distortions. Substrate connectivity was restored, the structure was optimized with YASARA, and the structure was inserted into the membrane as described above. The equilibrated structure is shown in Fig. 7B.

The GumI complex is less stable than GumH; although the reactive distance remains below 4.0 Å for over 100 ns, dissociation eventually occurs in all replicas, producing a bimodal distribution (Fig. 7B). The glucuronate orientation is maintained by stacking with Trp68, while its carboxyl group interacts with Arg179. In inverting GT-Bs, catalysis typically involves an acidic residue or histidine. Here, only a histidine is present near the reactive center, but it lies on a flexible loop connected to helix N*α*4 (Fig. 2A), making a stable catalytic role unlikely.

The lipidic tail remains localized between helices N*α*2 and N*α*3 (Fig. 2A), consistent with the apo and donor-bound simulations. In contrast to GumH, the conformational ensemble sampled by GumI shows a narrower distribution of inter-domain opening angles, as indicated by the projection onto the first (twisting) and third (bending) normal modes (Fig. 7B).

Overall, Boltz provides suitable starting conformations for the Gums ternary complex, although manual refinement is required. Starting from product-bound models helps constrain substrates into reactive poses. The tail-binding mode remains consistent with previous simulations, showing a structured clamp in GumH and a more open groove in GumI.

In the predicted GumI complex (Fig. 7B), no canonical catalytic base—Asp, Glu, or His—is positioned near the substrate [30]. In contrast, in GumH the substrate lies near Glu279 (Fig. 7A), part of the conserved EX_7_E motif of GT4 enzymes [29], supporting a binding pose compatible with the expected retaining mechanism [30].

## 3 Discussion

### 3.1 GumH and GumI resemble monotopic membrane proteins

We used Boltz to predict both apo conformations and protein–substrate complexes of the two GTs involved in xanthan gum biosynthesis, GumH and GumI. GumH belongs to the CAZy GT4 family, one of the best characterized and most heterogeneous GT families, with several members crystallized in apo, substrate-bound, and full-complex forms [31–33]. GumI belongs to the GT94 family and is the only experimentally characterized member of this family.

For both proteins, the predicted apo and complex structures consistently reproduce the same secondary-structure organization. The AlphaFold3-predicted fold shows a similar architecture (Fig. S12), indicating that the overall fold of the apo proteins is currently the most reliable model obtainable with generative AI methods.

In contrast, the relative orientation of the two domains is less certain, as shown by the distribution of inter-domain distances across apo predictions. This variability likely reflects conformational bias in structure prediction models, which favor states represented in their training sets [34, 35]. For GT4 enzymes, apo crystal structures are typically open, whereas substrate-bound forms are closed [36–38]. Consistent with this bias, GumH remains open in the apo predictions and adopts a more closed conformation upon substrate binding.

GumH has been reported as a cytosolic protein [4]. However, our simulations reveal a hydrophobic patch mainly formed by helix N-*α*4 (Fig. 2A) that partially inserts into the bilayer. This behavior agrees with OPM predictions [26] and is observed in the apo, donor-bound, and full-complex simulations. The helix contains three threonine residues that form intrahelical hydrogen bonds while exposing their methyl groups toward the membrane, increasing the hydrophobic character of the surface-facing side. Threonine and serine are generally weak helix formers in solution because their side chains interfere with backbone hydrogen bonding [22, 39]. In the membrane environment, however, hydrophobic interactions of the methyl groups and hydrogen bonding with phospholipid headgroups stabilize the helix. Thus, although this organization may not favor solution stability, it supports stable helix formation within the bilayer in our simulations.

Additional insight comes from simulations starting from a partially unfolded helix (Fig. S13), which does not refold efficiently in solution. Moreover, the loop connected to helix N*α*2 (Fig. 2A), part of the acceptor clamp, also interacts with the membrane during unbiased simulations. OPM does not identify this loop as membrane-interacting because the predicted starting conformation is suboptimal for membrane insertion.

These observations indicate that AI-based structure prediction may not fully capture the orientation of hydrophobic patches relative to the membrane, as membrane effects are not explicitly included. This limitation is particularly pronounced for the flexible loop, which tends to unfold in GumH simulations. The loop sequence (Fig. S4) suggests membrane interaction, as it contains both hydrophobic and charged residues. However, Boltz predicts a folded conformation in which Phe45 stacks beneath an adjacent loop. This arrangement is unstable in simulations, where Phe45 dissociates, leading to local destabilization. Despite high confidence scores, this region is likely mispredicted because membrane context was absent during structure generation.

Further insight into GumH membrane association arises from comparison with two GT4 members (Fig. 8A): PimA [40], a peripheral protein, and PglA [41], proposed to be a monotopic membrane protein. All three share a similar secondary-structure motif in the acceptor-binding domain. Helix N*α*4 (Fig. 2A) is conserved in PglA but disrupted in PimA by a proline residue. This structural similarity suggests that GumH resembles a stably membrane-associated GT4 more than a peripheral enzyme.

**Figure 8.**
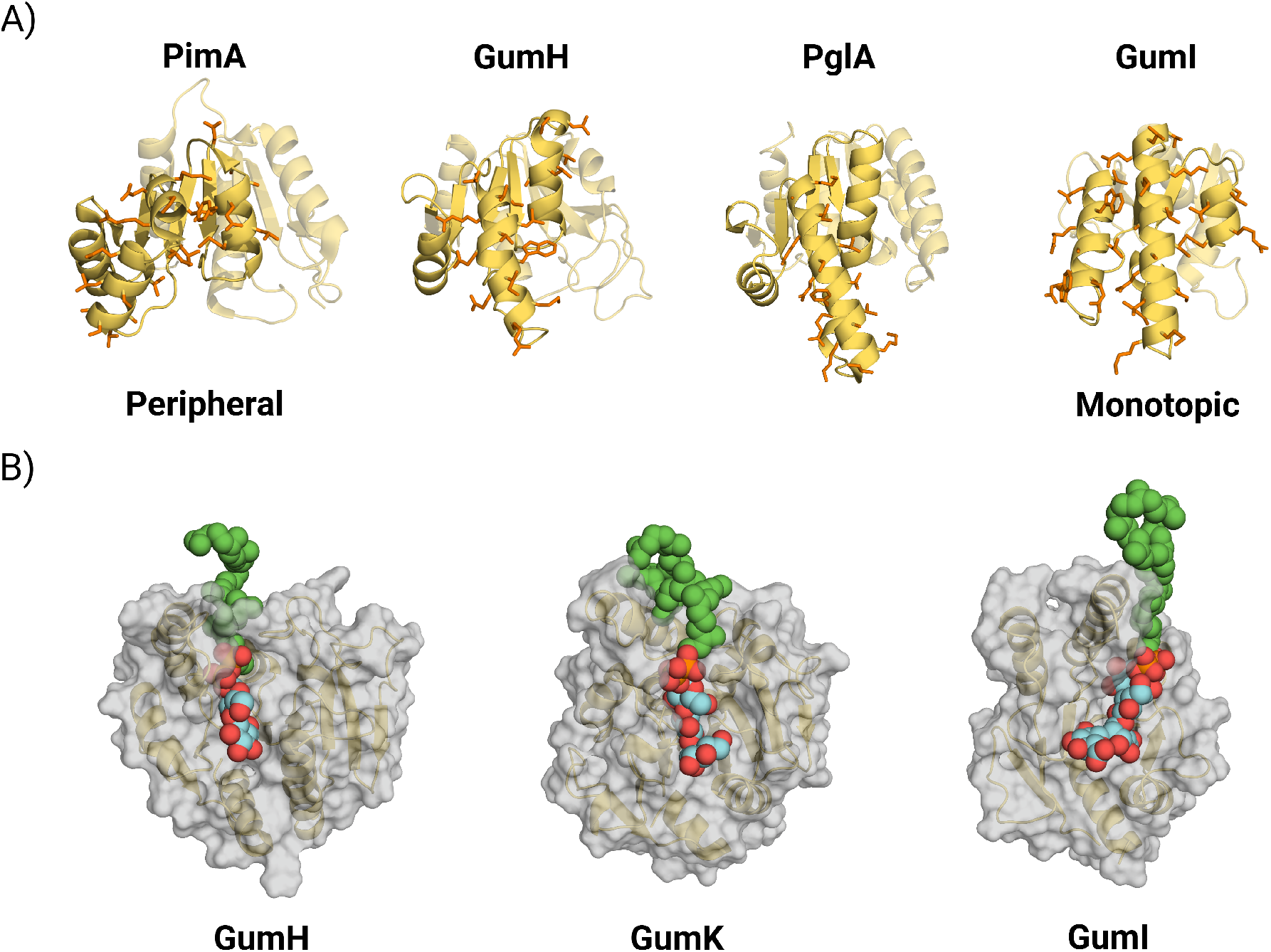
A) Comparison between the hydrophobic patches of the crystal structure of the peripheral GT4 protein PimA (PDB:2GEJ), the model0 of the apo form predicted in this work for GumH, the crystal structure of the monotopic GT4 PglA (PDB:8DQD), and model0 of the apo form predicted in this work for GumI. The motif highlighted in the picture is recurrent among GT4 members, and GumH has a structure similar to that of PglA, displaying a more complete hydrophobic helix than PimA. B) Comparison of the acceptor substrate interactions observed in this work for GumH and GumI with those previously reported for GumK. The localization of the tail-binding pocket on the acceptor domain appears to follow the same order as these enzymes in the biosynthetic pathway of xanthan gum.

Taken together, the reduced stability without a membrane, the composition and behavior of helix N*α*4, the instability of adjacent loops, and the similarity to membrane-anchored GT4 proteins, these findings indicate that GumH is unlikely to function as a purely cytosolic enzyme and is more stable when membrane-associated.

GumI instead displays features typical of a monotopic membrane protein. In particular, helix N*α*3 exposes two tryptophan residues (Fig. 2A), following the anchoring strategy observed in GumK [7]. Both enzymes are inverting GTs and share a similar membrane-interacting organization, characterized by tandem tryptophans on a hydrophobic helix.

### 3.2 GumI and GumH show different interdomain flexibility

As GT-Bs, GumI and GumH are composed of two domains that can undergo bending and twisting motions. These elementary motions are important for their activity, as they allow the binding of both soluble and hydrophobic substrates and enable the catalytic reaction. For GumK, we previously reported that the protein can open asymmetrically due to specific residues interacting at the interdomain interface [14].

GumH shows a stiffer opening motion when analyzed with ClustENMD than GumI. This behavior can be explained by the more structured linker region between the acceptor-binding domain and the donor-binding domain, which adopts an *α*-helical conformation instead of a flexible loop, as observed in GumI (Fig. 4B and D). GumI displays domain dynamics closer to those of GumK, as both have similar linker architectures that lead to overtwisted conformations during ClustENMD simulations (Fig. 1 and Fig. S8) [14].

The flexibility of the interdomain linker might be correlated with the tendency of GumI and GumH to open during unbiased MD simulations (Fig. 3C and D). Based on the conformational distributions sampled during the simulations, contrary to expectations, GumH, with a more rigid linker, tends to open more. In contrast, GumI keeps the two domains closer, and even when starting from model0, which we define as a semi-open state, it still samples the closed state in one of the three replicas.(Fig. 3D) . A complete loss of interdomain contacts in a protein with a flexible linker may lead to inactive states, such as overtwisted conformations. Since multiple conformations need to be explored, more time may be required for the domains to recover a closed state, potentially affecting catalytic efficiency. This correlation is consistent with previous results on GumK, in which the chemistry of the interdomain interface favored more closed conformations [14]. Similarly, our simulations suggest that the chemical composition of the interdomain interface in GumI, but not in GumH, helps preserve interdomain contacts and prevents fully open conformations that could negatively affect catalytic efficiency.

### 3.3 Differential donor substrate binding modes reflect different reaction mechanisms

Both GumI and GumH bind the same donor substrate, guanosine 5^*′*^-diphospho-*α*-D-mannose (GDP-Man), thereby facilitating comparison of their activities. The predicted structures of the donor substrate in complex with the enzymes reveal two distinct binding modes. In GumH, the main axis of the diphosphate group is oriented approximately perpendicular to the guanine plane, whereas in GumI, the two phosphate groups adopt a more parallel orientation (Fig. 6).

This difference in orientation may be explained by two key structural differences. GumH contains a longer helix facing the interdomain cleft (helix N*α*1). In the same structural position, the corresponding helix in GumI (helix N*α*1) is shorter and is not part of the interdomain cleft. In GumH, this helix favors the perpendicular orientation, as the negatively charged diphosphate group can be stabilized by the positive end of the helix dipole. Furthermore, GumH presents an apolar residue, Ala284, which, in the same structural position in GumI, is replaced by a hydrogen-bond donor, Thr260, thereby stabilizing the more horizontal orientation of the diphosphate group. The orientation of the diphosphate group might correlate with the catalytic mechanism since a similar binding mode has been reported in other inverting GTs [14, 42].

We therefore suggest that the chemical environment interacting with the diphosphate group determines stereoselectivity. However, it is important to note the limitations of this approach, particularly the bias that Boltz may introduce due to the availability of prior crystallographic information. In fact, the same diphosphate orientations are obtained in the ternary complexes of both substrates and products, highlighting a limitation of this generative AI approach.

The predicted complexes were not fully stable during the simulations; however, GumH adopted a more closed conformation than GumI in the presence of the donor substrate, whereas GumI tended to open, following a pathway similar to that observed in the apo simulations.

Our models indicate that binding the donor substrate affects both enzymes differently. Some members of the GT4 family [43–45] have been reported to show a stable closed conformation in the presence of the donor substrate. Our results (Fig. 6) showed that this applied to GumH, but not GumI, indicating differential substrate-binding effects. However, the bias introduced by Boltz in the starting conformation of GumI may have led to an initially closed state that was neither optimal nor stable during simulations. Another limitation is the force field used to represent the donor substrate, as it is well known that CHARMM36 [46], like other additive force fields [47, 48], may struggle to maintain stable protein– carbohydrate complexes.

### 3.4 The binding mode of the acceptor substrates has implications for enzymatic activity

In simulations of the full complexes and apo proteins, we identified regions that interact with the lipid carrier undecaprenyl diphosphate. GumH features a defined clamp between helices N*α*4 and N*α*3 (Fig. 2A) that opens in the absence of the acceptor, enabling lipid-tail binding. In contrast, GumI lacks a defined clamp; instead, the lipidic tail binds an open groove near helices N*α*3 and N*α*4, both in the apo state and in the full complex.

This difference likely reflects distinct strategies for acceptor recognition. GumH binds a disaccharide, and in the predicted acceptor site, no canonical sugar-binding residues are observed. GumI instead binds a tetrasaccharide and presents typical carbohydrate-interacting residues, including Trp68 and Tyr71. Thus, GumI may not require a tight clamp, as stacking of sugar units on aromatic residues provides stabilization. In this respect, GumH resembles GumK: in both cases, a clamp secures the lipidic carrier while the sugar-binding region orients the oligosaccharide into a reactive conformation.

The interaction network in GumI also explains its tendency to open in the absence of substrate. In the full complex, the acidic acceptor interacts with Arg179 in the donor-binding domain, promoting domain closure. This closed conformation persists throughout the simulations, even as the system relaxes from the Boltz starting structure (Fig. 6B and Fig. 7B).

For both enzymes, acceptor placement is predicted with lower confidence than donor binding, and geometry optimization with YASARA [49] was required to correct ring conformations. The limited stability observed in full-complex simulations likely reflects imperfect oligosaccharide docking. The prediction for GumI is likely less accurate than that for GumH because it involves a more complex substrate that lacks structural templates.

The lipid tail-binding region can nevertheless be defined with reasonable confidence, given the consistency of phospholipid positions in apo simulations and the predicted tail placement in the complexes. Comparison of this region in GumH, GumK, and GumI (Fig. 8B) reveals a progressive shift across the acceptor-binding domain. In GumH, the clamp is positioned on the left side (relative to Fig. 8B); in GumK it is more central; in GumI it is located on the right.

This shift likely has a geometric basis. In all three enzymes, the donor carbohydrate occupies the same position (on the left) within the active site. As the acceptor length increases along the pathway, the binding site shifts to the right, keeping the terminal reactive sugar near the donor. GumH binds the shortest acceptor, thereby positioning it close to the donor. GumK and GumI bind progressively longer substrates, requiring lateral displacement of the binding pocket to accommodate the growing oligosaccharide while preserving catalytic geometry.

An alternative possibility is that these enzymes interact at the membrane to exchange substrates, forming a multiprotein assembly, as proposed for a related system in *Rhizobium* [50]. However, no experimental evidence currently supports this model, and co-folding attempts did not yield a geometrically plausible complex.

### 3.5 The ternary complexes are consistent with two distinct enzymatic mechanisms

Despite the low confidence of the predicted GumH complex, which dissociates after 200–300 ns (Fig. S10D), the substrate orientation remains compatible with catalysis. The donor is positioned such that the phosphoacetal bond lies near the conserved Glu279–Lys203 pair (Fig. 6A and Fig. 7A), a feature common in GT4 enzymes [43, 51–53]. Retaining GTs such as GumH are proposed to follow an S_N_i mechanism [30], where spontaneous donor ionization generates an oxocarbenium ion and phosphate. The nucleophile then attacks from the same side as the leaving group. The Glu279–Lys203 pair may stabilize the developing negative charge in the transition state, thereby facilitating ionization.

The oxocarbenium intermediate is planar and, in principle, allows nucleophilic attack from either face. In the predicted complex (Fig. 7A), both C3 and C4 hydroxyl groups of the terminal acceptor unit are positioned near the donor anomeric carbon and could attack from opposite faces (C4 anti, C3 syn). However, simulations indicate that C3 is the preferred nucleophile because it forms a hydrogen bond with the *α*-phosphate group, thereby enhancing its nucleophilicity upon ionization of the donor. In AceA from *A. xylinum*, mutation of residues corresponding to Glu279 and Lys203 inactivates the enzyme [54], supporting their catalytic role. Overall, the predicted geometry is consistent with the experimentally observed regiospecificity of GumH.

The GumI complex is less stable during simulations (Fig. S10C), likely reflecting reduced accuracy of the initial prediction due to limited structural homologs in the training set. Nevertheless, stacking of the reactive *β*-glucuronate on Trp68 via its *α*-face defines a plausible reactive arrangement. Inverting GTs generally employ an Asp, Glu, or His residue to activate the acceptor hydroxyl in an S_N_2 mechanism. Here, no such residue is stably positioned near the reactive *α*-mannose–GDP and *β*-glucuronate pair. The only candidate, His112, lies on a flexible loop and is unlikely to act as a stable catalytic base.

The reactive oxygen in the acceptor (Fig. 7B) corresponds to a *β*-hydroxyl relative to the C5 carboxyl group of glucuronate. This geometry may exert dual effects. First, the nearby negative charge could interfere with a classical catalytic base. Second, the *β* configuration permits a transient intramolecular hydrogen bond between the hydroxyl and carboxyl group, potentially enhancing nucleophilicity and suggesting substrate-assisted catalysis. Additional support comes from Asn108, located near the reactive center; in other GT94 members, this position is occupied by Asp (UniProt: A0A430CIB1, A0A378RK98, A0A2I8AAR2, A0A2C9DCF8, A0A1Z4R3G0). This substitution suggests that GumI may lack a catalytic acid to achieve C4 regiospecificity on acidic substrates, whereas Asp-containing GT94 enzymes may target non-acidic substrates or exhibit alternative regiospecificity.

However, the intramolecular hydrogen bond between the carboxyl and *β*-hydroxyl is geometrically constrained by the ring, limiting the angle to 150°, below the ideal 180°. An alternative mechanism is concerted deprotonation of the *β*-hydroxyl by a phosphate group during nucleophilic attack. Substrate-assisted catalysis is common in retaining mechanisms but rare among inverting enzymes [30], with one example being AaNGT [42]. Nonetheless, inverting GT-B enzymes remain poorly characterized.

## 4 Conclusion

In this work, we have predicted conformations of GumH and GumI complexes that are chemically reasonable and compatible with catalysis. The combination of complementary computational methods with generative AI provides valuable qualitative insights into the activity of weakly characterized GTs. Further improvements, such as a better description of oligosaccharide interactions using more advanced force fields (e.g. polarizable models) and the use of AI-generated poses to guide more exploratory simulation approaches, could help to infer the activity of multiple members of poorly characterized GT families.

## 5 Methods

### 5.1 Structure prediction and system preparation

The structures of the GTs and their substrate-bound complexes were predicted using release 2.1.1 of Boltz [17]. Since all predicted models were subsequently used in atomistic molecular-dynamics simulations, the N- and C-terminal regions were excluded from the predictions to reduce the overall system size and avoid poorly predicted terminal segments. The amino acid sequences used for the predictions are listed in Table S1.

Donor substrates were specified in the Boltz input YAML files using their corresponding Chemical Component Dictionary (CCD) identifiers, whereas acceptor substrates were provided as SMILES strings with explicitly defined stereocenters to preserve carbohydrate chirality (Table S2). For product complexes, the initial substrate molecules were recovered by manually modifying bond connectivity in *PyMOL* [**pymol**], followed by energy minimization of the resulting complexes in vacuum using the standard optimization protocol implemented in *YASARA* [49].

The resulting protein–substrate complexes were oriented relative to a Gram-negative inner membrane using the OPM server [26]. System assembly, membrane construction, and solvation were performed using CHARMM-GUI[55], targeting an ionic strength of 0.15 M. The membrane lipid composition DYPE:DYPG:TYCL2 = 4:1:2.5 was selected to approximate as closely as possible the native inner-membrane environment of *Xanthomonas* species [28].

### 5.2 Normal mode analysis and ClustENMD calculations

Normal mode analysis (NMA) was performed using the Python library ProDy [24]. For each enzyme, GumI and GumH, two conformations were used as input: model0, the predicted apo form, and a reference closed conformation (conf0) extracted from the ternary complex prediction. Prior to the NMA, the proteins were oriented in a Gram-negative inner membrane bilayer using the OPM web server. A membrane elastic network model was subsequently built using the exANM[25] class implemented in ProDy. The membrane geometry followed the default settings: a face-centered cubic lattice with a node radius of 3.1 Å, no convex hull, and a membrane thickness of 25 Å, allowing proper insertion of the proteins’ membrane-associated regions. The protein’s elastic network model was constructed from only C_*α*_ atoms, and 20 normal modes were calculated for each system.

Conformational ensembles were generated using ClustENMD[27] class, employing the first three normal modes to define the deformation vectors. A target RMSD of 1 Å was applied at each generation, and a total of 30 generations were performed. The number of clusters was increased linearly at the end of each generation, with a step size of 10, starting from 10 clusters in the first generation. Each newly generated conformation was heated to 300 K and subsequently energy-minimized in implicit solvent using the default AMBER14 force field. Three simulations per protein were run to collect enough statistics on the conformations.

### 5.3 Force field parametrization

All protein, lipid, and carbohydrate components were described using the CHARMM36 force field[56]. The topology of the acceptor substrate was generated by merging the CHARMM carbohydrate parameters for the oligosaccharidic moiety with the CHARMM topology of undecaprenyl diphosphate, following established protocols for lipid-linked oligosaccharides [46]. The donor substrate was treated using a similar approach: the monophosphorylated sugar moiety was described using CHARMM carbohydrate parameters, whereas the guanosine monophosphate group was modeled using the CHARMM General Force Field (CGenFF) [57].

### 5.4 Equilibration protocol

All equilibration simulations were performed at 300 K using the velocity-rescaling thermostat [58], which was replaced by the Nosé–Hoover thermostat [59] during the final equilibration step. Pressure was maintained at 1 bar using the C-rescale barostat [60] during the initial equilibration phases and switched to the Parrinello–Rahman barostat [61] for the final equilibration.

Each system was equilibrated using a multi-step protocol consisting of two restrained NVT equilibration phases followed by six NPT equilibration steps, during which the force constant defining positional restraints was gradually decreased from 4000kJ mol^−1^ nm^−2^ to zero. These restraints were applied independently to protein backbone atoms, side chains, membrane lipids, and donor and acceptor substrates.

The final NPT equilibration step was performed differently for each system. For the apo protein systems of GumH and GumI, a 1 ns equilibration was carried out with a soft positional restraint of 50kJ mol^−1^ nm^−2^ applied to the backbone atoms. For the protein–donor complexes, a 10 ns equilibration was performed using a soft backbone restraint. For the full complexes, a total equilibration time of 100 ns was applied, during which backbone positional restraints were maintained, and an upper-wall restraint was imposed on the reactive distance between the nucleophilic hydroxyl group of the acceptor substrate and the anomeric carbon of the donor substrate. This restraint was implemented using PLUMED[62] in order to prevent divergence of the starting reactive complex.

### 5.5 Production simulations

Production simulations were initiated from the final equilibrated configurations. Pressure was maintained at 1 bar using a semi-isotropic Parrinello–Rahman barostat [61] with a coupling constant of 5.0 ps and a compressibility of 4.5 × 10^−5^ bar^−1^. Temperature was controlled at 300 K using the Nosé–Hoover thermostat [59] with a coupling constant of 2.0 ps.

For the apo proteins in solution, after an initial 10 ns unrestrained equilibration, production simulations of 1 µs were performed in triplicate. For the protein–donor substrate complexes, 500 ns production simulations were run in triplicate. Similarly, for the ternary complexes, 500 ns production simulations were performed in triplicate.

All simulations employed a cutoff of 1.2 nm for short-range van der Waals and electrostatic interactions, in accordance with CHARMM force field recommendations [56]. Long-range electrostatics were treated using the particle-mesh Ewald method, and standard long-range energy and pressure corrections were applied.

## Data Availability Statement

Data and scripts used for running and analyzing simulations, and for generating the figures, are available from https://github.com/gcourtade/papers/tree/master/2026/GumH_GumI

## Acknowledgements

G.C. gratefully acknowledges funding by the Novo Nordisk Foundation (grant number NNF22OC0073963 awarded to G.C.). This study made use of NMRbox: National Center for Biomolecular NMR Data Processing and Analysis, a Biomedical Technology Research Resource (BTRR), which is supported by NIH grant P41GM111135 (NIGMS).

## Conflict of Interest

The authors declare that they have no conflicts of interest.

## Notes

https://github.com/gcourtade/papers/tree/master/2026/GumH_GumI

### Competing Interest Statement

The authors have declared no competing interest.

### Summary of Updates

Figure 1 revised; grammatical and orthographic mistakes fixed

