## Supplementary Information for "Comparative study of two xanthan gum glycosyltransferases combining AI structure predictions and molecular modeling"

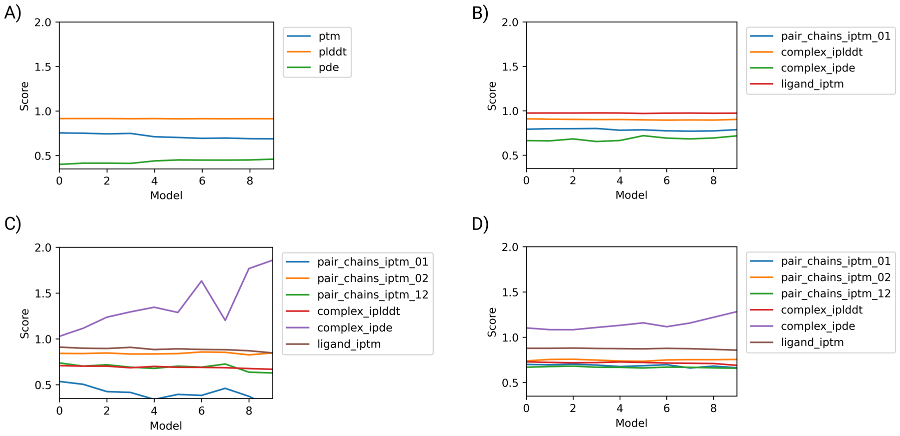


Figure S1: A) Cconfidence scores the ten models predicted using Boltz1 for the apo form of GumH. The pLDDT shows locally confident folding that is consistent among the ten models, indicating good reproducibility. The pDE and pTM show differences across models, suggesting intrinsic flexibility in the protein. Overall, Boltz-1 consistently predicts a reliable fold for GumH, as indicated by high, reproducible local confidence scores and a stable global topology across independent models. B) Confidence scores for the GumH–donor complex, showing a similar behavior compared to the apo protein prediction. However, the overall complex displays lower confidence, particularly at the interface. C) Confidence scores of the ternary complex of GumH with the two substrates. The ten models are consistent in local atomic positions, but relative positioning within the complex shows very low confidence, which increases slightly in some models and is mainly associated with the lipid tail. D) Confidence scores of the ternary complex with the products. In this case, the overall prediction quality is higher; however, the relative atomic positions at the complex interface remain critical.

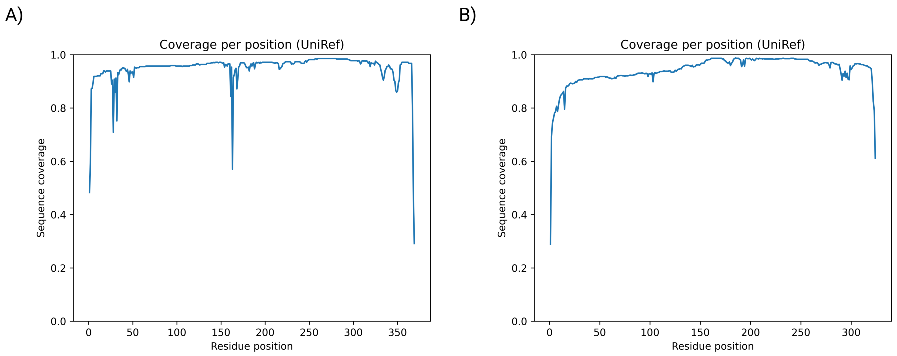


Figure S2: A) Coverage sequence of GumH in the prediction of the apo form using Boltz1. B) Sequence coverage of GumI in the apo form during Boltz1 prediction.

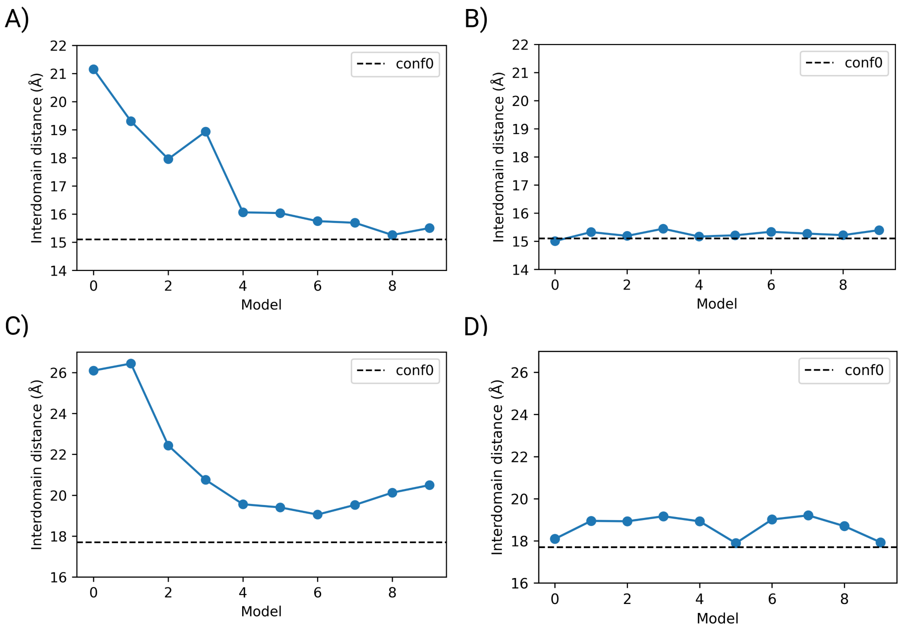


Figure S3: A) Interdomain distance among the ten predicted models of the apo form of GumH. The dashed line refers to the reference closed conformation of GumH (conf0). Variability in the degree of opening is observed, with model0 appearing the most open. B) Interdomain distance for the ten models predicted for the GumH–donor complex, which show a more consistent closed conformation compared to the reference closed structure (conf0). C) Interdomain distance of the GumI apo form predicted with Boltz1. The 10 generated models exhibit similar behavior, with the protein adopting a more open conformation than the reference closed state (conf0), as indicated by the dashed line. D) Interdomain distance of the GumI–donor complex predicted using Boltz1, showing that the ten folds are similar to the reference closed conformation (conf0) in terms of degree of opening.

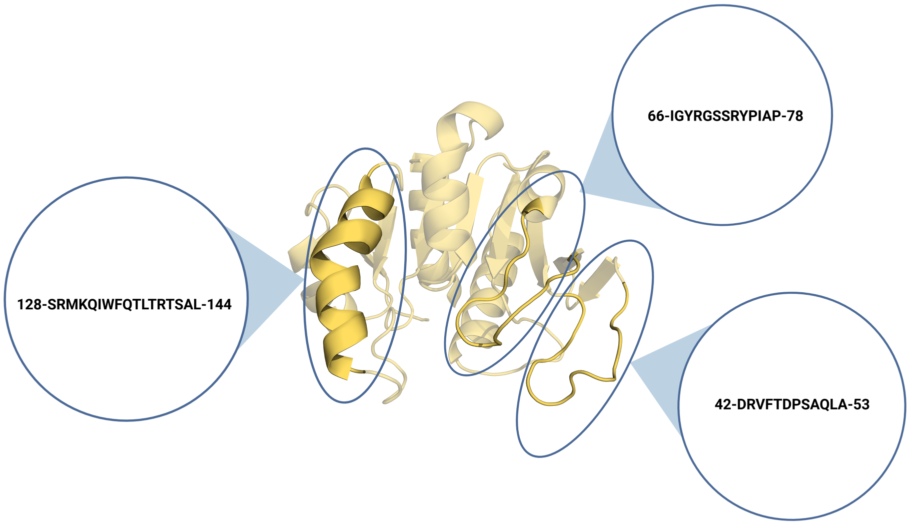


Figure S4: Sequences of the hydrophobic regions of GumH.

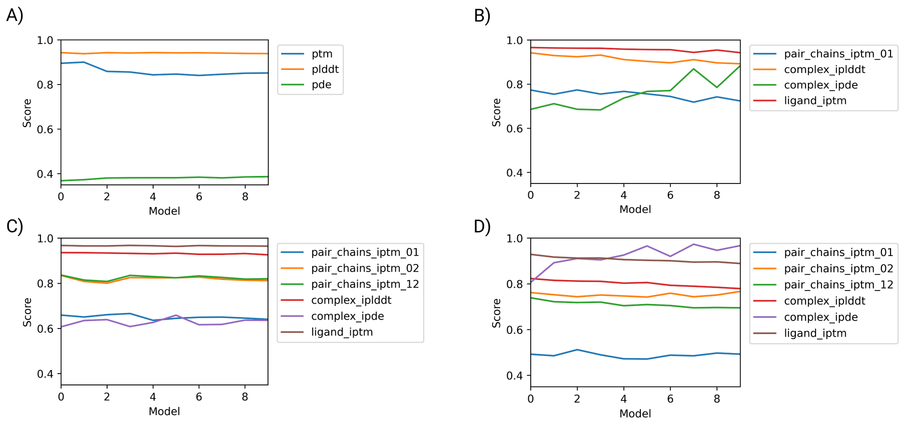


Figure S5: A) Confidence scores of the apo form of GumI, showing high confidence in the local fold but lower confidence in the overall fold, probably due to the relative positioning of the two domains. B) Confidence scores of the ten predictions for the GumI–donor complex. The models are consistent in their confidence, but the relative positioning of the ligand atoms with respect to the protein increases in the later models. C) Confidence scores of the ternary complex with GumI binding the two substrates. The ten models are consistent in their confidence estimates, and the greater consistency of ipDE is likely due to the absence of a lipid tail on the acceptor substrate. D) Confidence scores of the prediction of GumI in complex with the products. The confidence in the relative positioning of the ligand atoms with respect to the protein is lower than in the previous case.

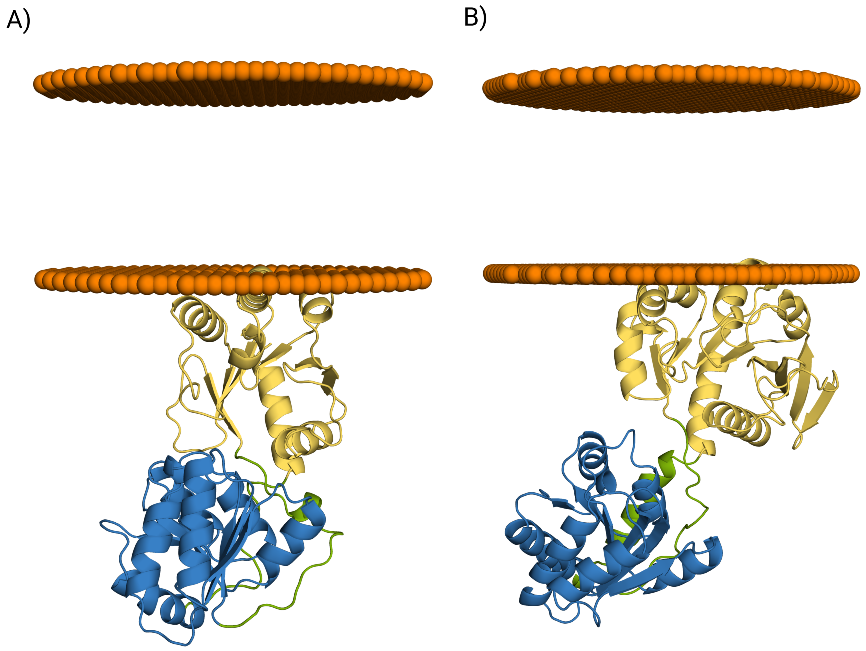


Figure S6: A) OPM-oriented GumI in an inner-membrane Gram-negative bacterium. B) GumH oriented in an inner-membrane Gram-negative bacteria using OPM.

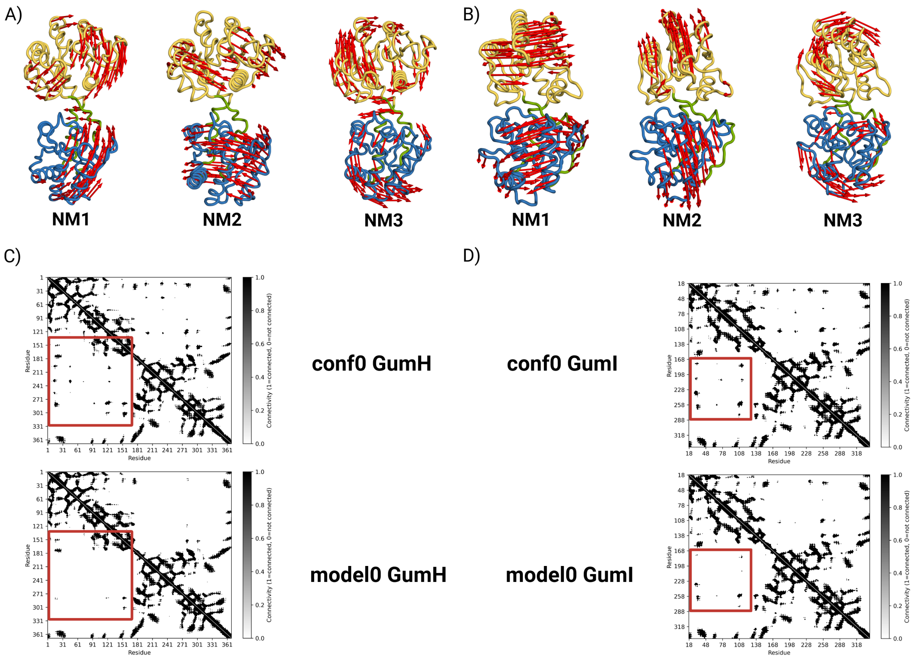


Figure S7: A) First three normal modes of model0 for the apo form of GumH predicted using Boltz1. B) First three normal modes computed for model0 of the apo form prediction of GumI. C) Comparison of the Kirchhoff matrices showing the different topology of the elastic network for conf0 and model0 of the apo form prediction of GumH. The former is considered a closed conformation, and the red boxes highlight the loss of interdomain contacts in model0 respect to conf0, which explains the large difference observed in the normal mode solutions between the two conformations. D) Kirchhoff matrices of conf0 and model0 of the apo form prediction of GumI. In this case, conf0 represents a more closed conformation; however, compared to GumH, more interdomain contacts remain in model0, as shown by the red boxes. This is the reasion of the inversion in the order of the normal modes when comparing conf0 with model0.

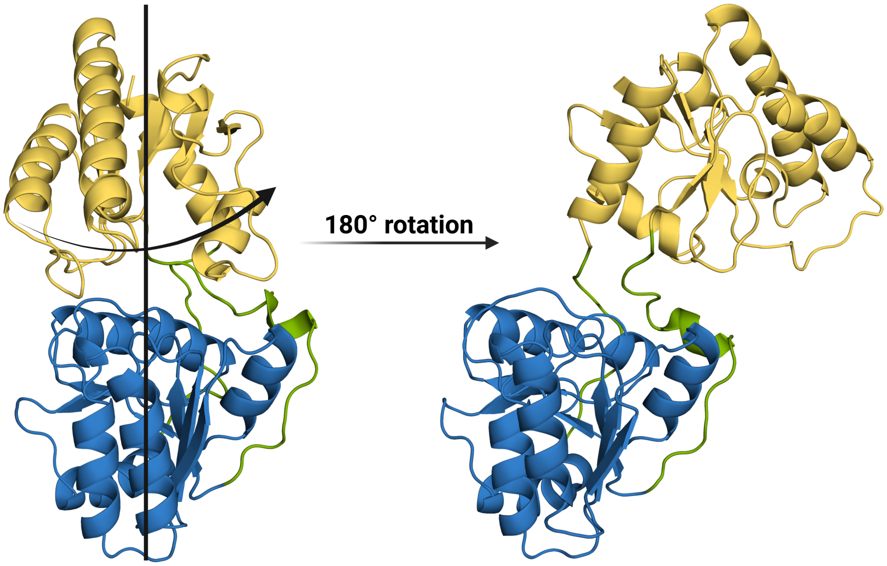


Figure S8: On the right, GumI conf0, ont the left an example of overtwisted conformation sampled during ClustENMD simulation using conf0 as starting confromation . The aceptor binding domain is 180° rotated around the main axis as shown by the arrow on the conf0.

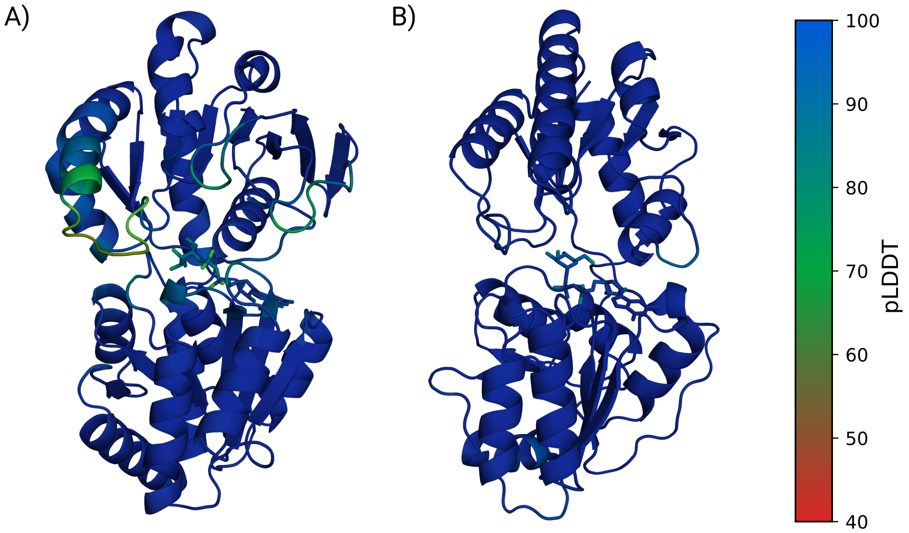


Figure S9: A) pLDDT of model0 for GumH-donor complex. B) pLDDT of model0 of GumI-donor complex.

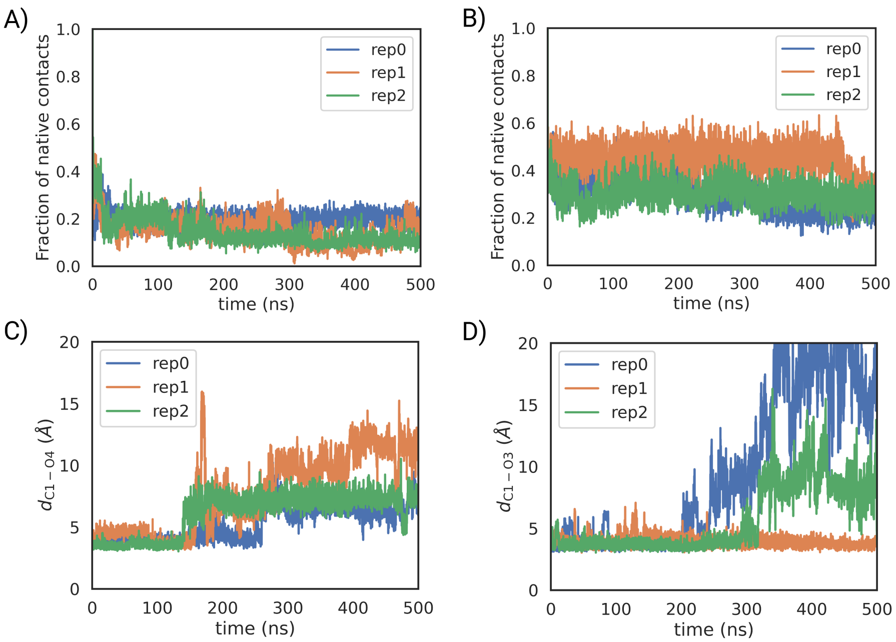


Figure S10: A) Fraction of native contacts during the molecular simulation of model0 predicted for the GumI–donor complex. B) Fraction of native contacts during the molecular simulation of model0 predicted for the GumH–donor complex. C) Reactive distance between the anomeric carbon of the donor substrate and the reactive oxygen of the acceptor substrate in the ternary complex of GumI during the unbiased molecular simulation of the model0 predicted conformation. D) Reactive distance between the anomeric carbon of the donor substrate and the reactive oxygen of the acceptor substrate in the ternary complex of GumH during the molecular simulation of the model2 predicted conformation.

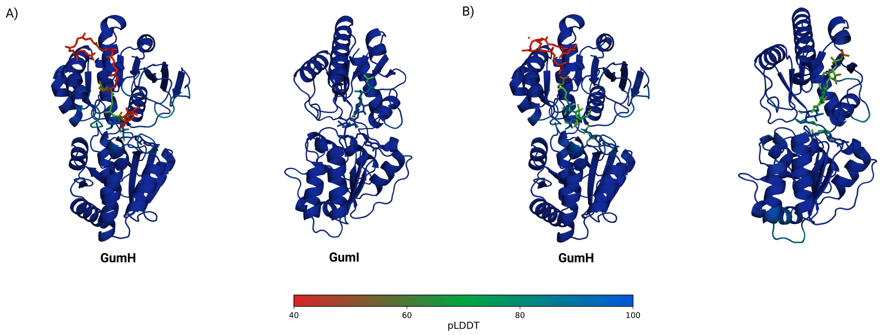


Figure S11: A) pLDDT score of model0 for the ternary complexes of GumH and GUmI in complex with the substrates. pLDDT score of model2 and model0 for the ternary complexes, respectively, for GumH and GumI in complex with the products.

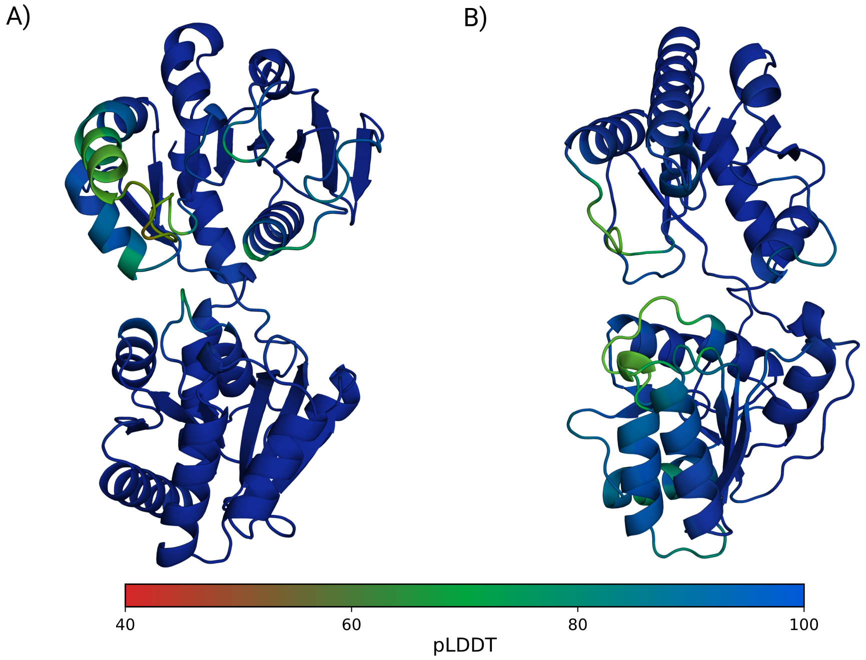


Figure S12: A) model0 of the apo form of GumH was predicted using the AlphaFold3 web server. B) model0 of the apo form of GumI predicted using the AlphaFold3 web server.

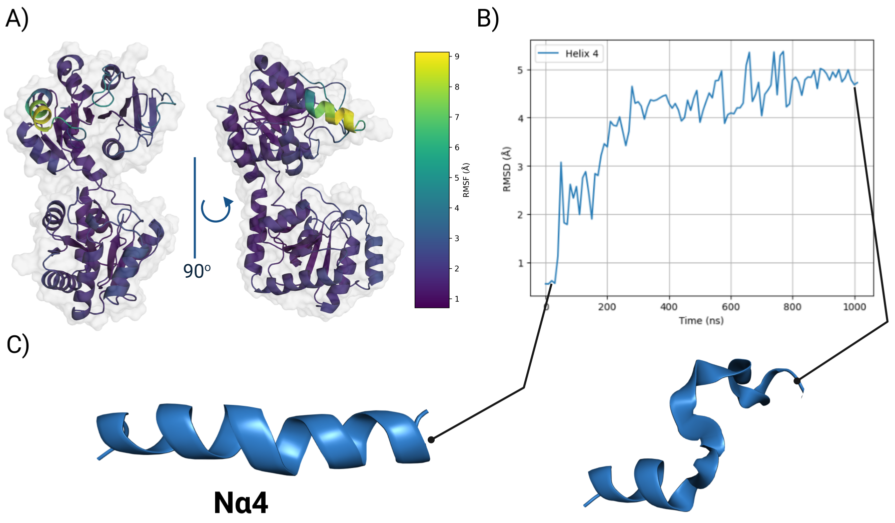


Figure S13: A) RMSF of the best-scored AlphaFold-predicted structure of GumH computed over 1 µs of unbiased molecular dynamics simulation in aqueous solution without a membrane. B) RMSD of the hydrophobic helix N$\alpha$4 during the aqueous simulation. C) Representative conformations of the hydrophobic helix at the beginning and at the end of the simulation. The helix displays structural instability when simulated in solution in the absence of a membrane.

Table S1: Input sequences for Boltz1 prediction

| **Protein** | **Sequence** |
|---|---|
| GumH | MKVVHVVRQFHPSIGGMEEVVLNVARQHQANSADTVEIVTLDRVFTD PSAQLAQHELHQGLSITRIGYRGSSRYPIAPSVLGAIRSADVVHLHGIDF FYDYLALTKPLHGKPMVVSTHGGFFHTAYASRMKQIWFQTLTRTSA LAYARVIATSENDGDLFAKVVAPSRLRVIENGVDVEKYAGQGARAPG RTMLYFGRWSVNKGLIETLELLQAALTRDPQWRLIIAGREYDLNEAD LRKAIAERGLQDKVQLSMSPSQQQLCALMQQAQFFVCLSRHEGFGIAA VEAMSAGLIPILSDIPPFVRLATESGQGVIVNRDRIQAAADSVQALALQ ANADFDARRTATMAYVARYDWRHVVGRYIDEYHAA |
| GumI | ITVLFSTEKPNANTNPYLTQLYDALPDAVQPRFFSMREALLSRYDVLH LHWPEYLLRHPSKMGTLAKQACAALLLMKLQLTGTPVVRTLHNLAP HEDRGWRERALLRWIDQLTRRWIRINATTPVRPPFTDTILHGHYRDW FATMEQSTTLPGRLLHFGLIRPYKGVEVLLDVMRDVQDPRLSLRIVGN PATPQMRTLVETACAQDARISALLAYVEEPVLAREVSACELVVLPYK QMHNSGTLLLALSLARPVLAPWSESNAAIADEVGPGWVFLYEGEFDA ALLSGMLDQVRAAPRGPAPDLSQRDWPRIGQLHYRTYLEAL |

Table S2: SMILES codes used as input in Boltz-1 for the prediction of the full donor–acceptor complexes.

| Substrate | SMILES code |
|---|---|
| GumH_acceptor | C/C(C)=C\CC/C(C)=C/CC/C(C)=C/CC/C(C)=C\CC/C(C)=C\CC/C(C)=C\CC/C(C)=C\CC/C(C)=C\CC/C(C)=C\CC/C(C)=C\CC/C(C)=C\CC/C(C)=C\COP(=O)([O-])OP(=O)([O-])O[C@H]3O[C@H](CO)[C@@H](O[C@@H]2O[C@H](CO)[C@@H](O)[C@H](O[C@H]1O[C@H](CO)[C@@H](O)[C@H](O)[C@@H]1O)[C@H]2O)[C@H](O)[C@H]3O |
| GumI_acceptor | O=C([O-])[C@H]4O[C@@H](O[C@H]1[C@@H](O)[C@H](O)[C@@H](CO)O[C@@H]1O[C@H]3[C@H](O)[C@@H](CO)O[C@@H](O[C@H]2[C@H](O)[C@@H](O)[C@@H](OP(=O)([O-])OP(=O)([O-])[O-])O[C@@H]2CO)[C@@H]3O)[C@H](O)[C@@H](O)[C@@H]4O[C@@H]5O[C@H](CO)[C@@H](O)[C@H](O)[C@@H]5O |
